# Impact of mating strategies on life-history traits in the alien land snail *Rumina decollata*

**DOI:** 10.1101/2025.01.31.635896

**Authors:** Julia Pizá, Lara Cifola, Melisa Perl, Matías Abafatori, Nicolás Bonel

## Abstract

Climate change and global transport are driving species introductions worldwide, leading to economic and ecological consequences. Hermaphroditic organisms are able to reproduce with any conspecific, and some can self-fertilize, enhancing their potential for population establishment despite low initial densities in colonization events. This study examines how mating strategies influence the life history traits of the alien land snail *Rumina decollata* by comparing individuals subjected to facultative cross-fertilization or enforced self-fertilization over two laboratory-reared generations. Key life history traits—including size and age at first reproduction, fecundity, hatching time, and juvenile survival—were measured, alongside individual growth and shell morphometry. Self-fertilizing individuals exhibited higher body weight at first clutch but lower fecundity and delayed reproduction compared to cross-fertilizers. Selfing offspring (F_2_) took longer to hatch and had lower survival rates, indicating significant inbreeding depression. Selfed snails of F_1_ grew faster than outcrossers, but experienced a decline in growth in F_2_, consistent with inbreeding depression. Conversely, shell shape remained similar between mating treatments. Although selfing imposed fitness costs, 32% of self-fertilizing individuals produced viable offspring, highlighting their ability to establish in new environments. This study improves our understanding of how *R. decollata*’s reproductive strategies shape life history traits under environmental constraints.

**SIMPLE SUMMARY:** Species are spreading to new regions due to climate change and global transport, often causing environmental and economic challenges. Hermaphrodite species that can reproduce by cross or self-fertilization can establish populations even when few individuals are present. In our study, we examined how these two reproductive strategies affect a land snail’s growth, reproduction, and survival. We found that self-fertilizing snails grew larger and heavier but laid fewer eggs and reproduced later than those that mated with others. Additionally, their offspring took longer to hatch and had lower survival rates. These results suggest that self-fertilization comes with costs, likely due to inbreeding depression, but it still allows the species to establish in new environments. Despite lower survival, one-third of the self-fertilizing snails produced viable offspring, showing that this strategy can support population growth when mates are scarce. As this species continues to expand into new areas, our findings help explain how it adapts to different environments. Understanding these reproductive strategies is important for predicting its spread and managing its potential impact on ecosystems.

## Introduction

Climate change and global transport are contributing to the introduction of species around the world, with associated economic and ecological impacts [1,2]. Colonization events are often initiated by a limited number of individuals, resulting in low population densities [3,4]. Under such conditions, the reduced probability of finding a mate can result in decreased reproductive success [5]. It is essential, then, to investigate how species respond to environmental constraints, such as low mate availability.

Unlike species with separate sexes, hermaphroditic organisms benefit from the ability to reproduce with any conspecific, which enhances their potential to establish populations and surmount challenges posed by low population densities during the early stages of colonization. Moreover, some hermaphroditic species can self-fertilize when mates are unavailable, a reproductive assurance strategy that ensures reproductive success under such conditions [6–9]. The mating systems exhibited by hermaphroditic species in natural populations can vary and are generally categorized as: (i) predominantly self-fertilizing (selfers), (ii) predominantly cross-fertilizing (outcrossers), or (iii) mixed [10].

A species’ ‘preferential mating system’ can be defined by its selfing rates under ideal conditions, free from environmental constraints such as mate scarcity. While many species are thought to exhibit mixed mating systems, they may actually prefer outcrossing but resort to self-fertilization in response to environmental constraints [6,8]. When mates are unavailable, preferential outcrossers face two options: self-fertilize, incurring the costs of inbreeding depression, or delay reproduction while waiting for a mate, risking mortality before reproducing. Self-fertilization increases homozygosity within individuals, which can result in inbreeding depression—manifesting as reduced viability, fertility, or even complete failure to develop in selfed offspring [10–12]. For preferential outcrossers, isolated individuals are expected to delay their age at first reproduction (i.e., waiting time) when mates are unavailable. In contrast, preferential selfers are expected to self-fertilize as soon as they reach sexual maturity, regardless of mate availability [6].

The cosmopolitan land snail *Rumina decollata* (Linnaeus, 1758) is a simultaneously hermaphroditic that exhibits a mixed mating system [13]. Native to the Mediterranean region, *R*. *decollata* has been widely introduced to Asia and the Americas through human activities. In Argentina, its presence was first reported in 1988 in Buenos Aires [14], and since then, it has steadily expanded its range across the country—from Patagonia to its northernmost regions—demonstrating an exceptional ability to colonize diverse ecosystems with varying environmental conditions [15]. Previous studies on allozyme data and lab experiments suggested that self-fertilization is a regular, if not the predominant, mode of reproduction in this species [16]. This hypothesis is supported by a high percentage of isolated snails observed to self-fertilize and the low genetic variability reported in native and introduced populations based on alloenzymes and microsatellites analyses [16–19]. However, a more comprehensive study using nine microsatellites revealed significant allelic variation and heterozygous genotypes, suggesting that cross-fertilization rates in *R*. *decollata* are higher than previously estimated [20]. These findings challenge the notion that self-fertilization is the predominant reproductive strategy in this species. Despite its remarkable colonization success, the extent to which its mating system strategy influences key life history traits remains poorly understood. Addressing this gap is critical for uncovering the mechanisms driving *R*. *decollata*’s sustained range expansion and its ability to thrive in diverse and changing environments.

In this study, we investigated how mating strategies influence the life history traits of *R*. *decollata*, We established two experimental lines consisting of first- and second-generation laboratory-reared individuals under controlled conditions, each subjected to one of two mating treatments: facultative cross-fertilization (outcrossing treatment) and enforced self-fertilization (selfing treatment). For each treatment, we measured key life history traits, including size and age at first reproduction, fecundity, hatching time, and juvenile survival. We also estimated and compare individual growth and shell morphometrics.

If *R*. *decollata* is a preferential outcrosser that resorts to self-fertilization only when mates are unavailable, we hypothesize that enforced self-fertilization may also heighten inbreeding depression, resulting in lower fecundity and reduced juvenile survival in the selfing treatment. We also expect that selfing individuals will exhibit delayed egg laying (waiting time), leading to larger size and older age at first reproduction. This delay may enable selfing individuals to reallocate resources initially intended for reproduction toward growth, consistent with a growth-focused strategy [21]. If this resource allocation trade-off holds true, we further hypothesize that selfing snails will exhibit higher individual growth rates compared to outcrossed ones. Because shell growth is closely linked to energy allocation strategies, differences in growth rates may influence shell shape. However, the relationship between growth rate and shell morphology varies among gastropods, with faster growth sometimes leading to more elongated shells and, in other cases, resulting in more globular forms with lower spires [22,23]. Given this variability, we hypothesize that selfing snails, through reallocation of resources toward growth, will exhibit distinct shell shapes compared to their outcrossed counterparts. Whether selfing individuals develop more elongated or more globular shells may depend on the specific trade-offs in energy allocation between somatic growth and reproduction. These potential morphological differences could reflect adaptive responses to the reproductive constraints imposed by self-fertilization.

## Methods

### Study organism and experimental conditions

*Rumina decollata* (Stylommatophora: Achatinidae) is a terrestrial gastropod characterized by its elongated, cylindrical shell and distinctive truncated apex, which results from the natural loss of the initial shell whorls during decollation. It is a simultaneously hermaphroditic species that frequently engages in reciprocal copulation but also has the capacity for facultative self-fertilization [24,25]. The species reaches reproductive maturity at approximately two months and is iteroparous, laying multiple clutches of eggs throughout its lifespan [17]. Laboratory studies have reported variations in clutch size and incubation period depending on temperature conditions. On average, clutch size remains consistent at around 12 eggs [17], while incubation period varies with temperature—ranging from 30 days at 20°C [24] to longer durations at lower temperatures ([1]: 32 days at 22°C, 45 days at 18°C).

Throughout the experiment––and also during the establishment of experimental lines––snails were maintained under standard laboratory conditions (23±1 °C, a 16:8 photoperiod, and *ad*-*libitum* food in the form of ground lettuce). Individuals were kept in a 200 mL plastic box filled with a 3 cm thick layer of soil enriched with calcium carbonate, except when mass mating where they were put in a terrarium (38 x 28 x 8.5 cm) in order to facilitate mate encountering.

### Snail population sampled and establishment of experimental lines

#### Field sampling and breeding design

The first step was to collect snails from a wild population at the Universidad del Sur (38°42’4.32”S, 62°16’7.06”O) in Bahía Blanca (Argentina) on March 2022 and transported them to the lab where they were kept in controlled laboratory conditions (see above). After two weeks of acclimation to laboratory conditions, all individuals (n = 76) were grouped in a terrarium in which they freely mated for one week (mass mating). After mass mating, individuals were isolated in a box and eggs were collected immediately after we detected the first clutch (hereafter Group A). Once snails laid the first clutch, we transferred them to a new box to collect the second clutch (hereafter Group B). The offspring from each clutch were laid by the same mother, meaning that they belong to the same family and that they are full-sibs. These juveniles constituted the first generation of snails raised in controlled laboratory conditions (F_1_). One juvenile per clutch (hereafter focal snail) was raised and isolated before maturity, as previously mentioned, and kept until fully mature and ready for mating. We used these focal snails to generate two independently laboratory populations, each comprising approximately 40 to 70 reproductive individuals.

The experimental populations encompass two mating treatments: facultative cross-fertilization (hereafter ‘outcrossing’) and obligate self-fertilization (herafter ‘selfing’) each one corresponding to Group A and B, respectively. The outcrossing treatment consisted in pooling all virgin individuals (n = 70)–– kept in isolation until they were fully mature––in a terrarium in which they freely mated for one week (mass mating) to promote cross-fertilization. Given the reciprocal copulation in this species [24,25], each snail was likely to play both male and female roles many times during this period. After mass mating, individuals were re-isolated to lay eggs. In the selfing line, self-fertilization was enforced by keeping mature individuals isolated until they lay eggs (see Figure S1 in Supp. Inf. for more details on the breeding protocol).

### Assessing and comparing life history traits

For both mating treatments, we weighed each F_1_ snail when they laid their first clutch (laying date). Additionally, we determined the age (days) at first reproduction in order to assess whether snails in the selfing treatment exhibit a delay in egg laying, the so-called waiting time [6,10]. Eggs were counted in all F_2_ clutches right after collection, representing female fecundity under self- and cross-fertilization conditions. Hatching time was assessed as the number of days elapsed since egg laying until egg hatching. Once hatched, we counted hatchlings (F_2_ snails) in the boxes 14 days later and juvenile survival was estimated as the ratio of live juveniles over the number of eggs in each box. Then, these offspring were raised in isolation until maturity and became the parents of the next generation (F_3_ snails), on which we also evaluated life history traits (see Figure S1 in Supp. Inf. for more details on the breeding protocol).

We estimated apparent inbreeding depression (*AID*) as a proxy to evaluate differences in juvenile survival between mating treatments. Following the definition by Escobar et al. (2011), AID was calculated as: *AID* = 1 − *W_S_*/*W_O_*, where *W_S_* and *W_O_* stand for the mean juvenile survival of offspring from the selfing and the outcrossing treatments, respectively. The maximum inbreeding depression (*ID_max_*) was defined as 1 − *W_S_*, providing an upper limit to inbreeding depression in the selfing treatment, as survival cannot exceed 1 [26]. We focused on juvenile survival because inbreeding depression tends to be most pronounced at this stage of the life cycle [27,28].

### Individual growth rates and parameters

To assess individual growth variation between linetypes, we took photographs of each individuals once a month for the F_1_ individuals and every two weeks for the F_2_ snails. For both generations, we measured shell length using the ImageJ software (see Figure S2 in Supp. Inf. for specific details on the methodology). Then, we fitted the von Bertalanffy growth function with seasonal oscillations (hereafter SVBGF) [29–31]:

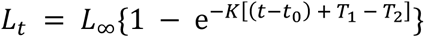

where

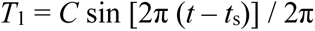

and

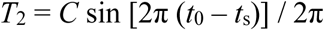

and where Lt is the predicted shell length at age t, L∞ is the asymptotic shell length, K is the growth constant of dimension time (year-1 in most seasonally oscillating growth curves) expressing the rate at which L∞ is approached, t0 is the theoretical age of the clam at shell length zero, C represents the relative amplitude of the seasonal oscillation and varies between 0 and 1 (0 indicating lack of summer–winter differences in growth, it was constrained to be ≤1 during model fitting), and ts is the starting point of the oscillation. When preliminary results of the SVBGF failed to converge to estimate the asymptotic shell length (L∞), we used the maximum shell length (L_max_) observed or reported at each study site to calculate the asymptotic shell length. To do this, we followed the equation suggested by Taylor [32]:

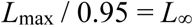

These values were fixed when we performed a second fit. Because the negative correlation between growth parameters (*K* and *L_∞_*) prevents making comparisons based on individual parameters [33,34], we used the growth performance index (GPI), which reflects the growth rate of a given organism of unit length. In other words, GPI can be viewed as the (theoretical) value of *K* that would occur in organisms with an L∞ value of 1 unit of length [35], and defined by Pauly and Munro [36] as:

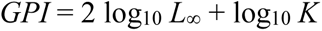

We calculated the GPI values for each mating treatment and generation based on growth parameters (*L_∞_* and *K*) obtained after fitting the SVBGF.

### Geometric morphometric analyses

To evaluate whether individual growth variation influenced shell morphology in *Rumina decollata*, we analyzed the final set of images obtained after one year of growth monitoring for each mating treatment (N = 40 per treatment). Images were digitized and processed using *tpsUtil* software (v. 1.82), while landmarks were placed on shell using *tpsDig2* software. A total of 13 landmarks were defined for the analysis, following established protocols [37].

To examine shell shape variation, we used *MorphoJ* software (version 1.08.01; [38]). The first step involved a *Procrustes* superimposition, which standardize the data by removing differences in position, orientation, and scale while preserving shape variation. This process also allowed for the calculation of centroid size, a proxy for overall shell size [37]. We then conducted a regression analysis between shape and centroid size, assessing significance through a permutation test to determine the presence of allometry—the relationship between shape changes and size differences [39]. To account for this effect, we used residuals from the regression for further analyses.

A covariance matrix of the average shell shape was constructed before performing a Principal Component Analysis (PCA) to visualize major axes of shape variation across populations [40]. Finally, a Canonical Variate Analysis (CVA) was conducted to statistically assess and enhance visualization of shape differences among groups. CVA identifies shape characteristics that best differentiate groups while minimizing within-group variation [41].

### Statistical analyses

All statistical analyses were performed in R 4.4.1. Linear models (LMs) were used for Gaussian continuous variables (e.g., body weight), while generalized linear models (GLMs) were implemented for Poisson-distributed variables (e.g., age at first reproduction, fecundity, hatching time). To account for overdispersion in Poisson models, identified as a ratio of residual deviance to residual degrees of freedom > 1.2, quasi-Poisson models were employed to address extra-Poisson variation. Generalized linear mixed models (GLMMs) were used for binomial variables (e.g., juvenile survival), implemented with the *lme4* package. Overdispersion in the binomial model was handled by including observation number as a random factor, following [42]. Model fit and treatment effects were assessed using *F*-tests for LMs and quasi-Poisson models. For GLMMs, fixed effects were tested using Likelihood-Ratio Tests (LRT), which compared models with and without the fixed factor of interest, while retaining the random effects structure. Random effects were evaluated using LRT, applying corrections as described in Zuur et al [43]; Chapter 5, pp. 123–125).

## Results

### Life history traits

Body weight of the F_1_ at first reproduction varied between treatments. The selfing treatment had the highest mean, with an increase of 21% in weight compared to the outcrossing treatment (Table 1, Figure 1A). The age at first reproduction had no significant effect on body weight when included as a covariate in the model to test the effect of mating treatments (*P* = 0.304; Table S1). However, when analyzed independently, snails from the selfing treatment laid their first clutch 39 days later than those from the outcrossing treatment (Table 1, Figure 1B). We found a significant effect of mating treatment on the female fecundity (Table 1, Figure 1C), the selfing snails laid 51% fewer eggs per individual than the outcrossed ones. When added as a covariate, body weight showed no effect on fecundity (*P* = 0.727; Table S1). On average, eggs from the selfing treatment hatched 4 days later than eggs from the outcrossing treatment (Table 1, Figure 1D). Juvenile survival of the F_2_ significantly differed between mating treatments. The survival of the selfing treatment (0.30) was lower compared to the outcrossing treatment (0.70; Table 1, Figure 1E). The apparent inbreeding depression (*AID*) for juvenile survival was estimated at 0.57. This indicated a 57% reduction in juvenile survival under selfing compared to outcrossing. The maximum inbreeding depression (*ID_max_*) was 0.70 (70%), which indicated the upper limit to the potential inbreeding depression observed in the selfing treatment.

**Figure 1.**
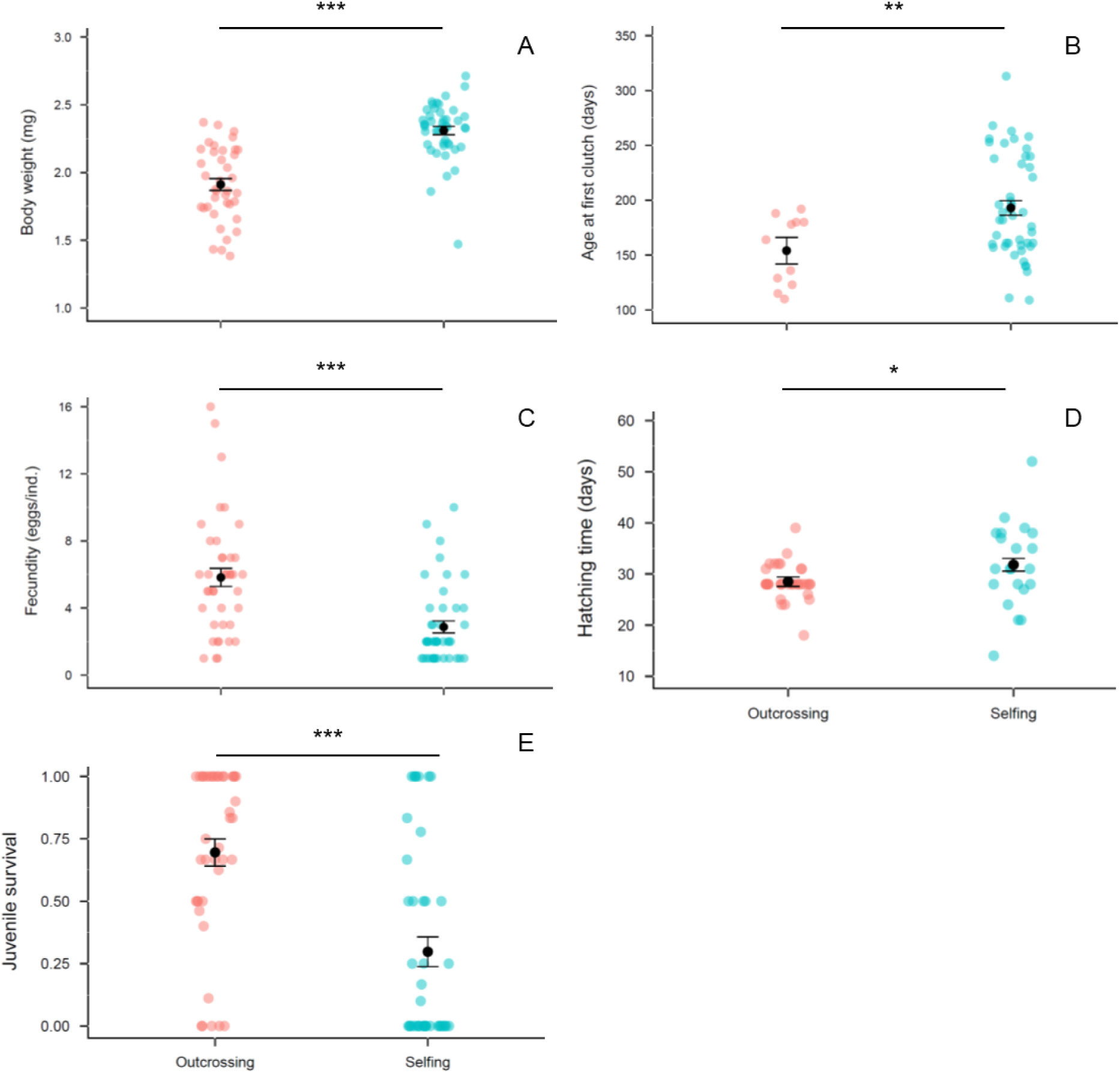
Fitness traits measured on F_1_ individuals of the cosmopolitan land snail *Rumina decollata*. A. Body weight at first reproduction. B. Age at first clutch (days). C. Fecundity, measured as the number of eggs per clutch per individual. D. Hatching time, the number of days elapsed from egg laying to hatching. E. Juvenile survival, proportion of eggs that become juveniles alive 14 days post-hatching. ∗*P* < 0.05; ∗∗*P* < 0.01; ∗∗∗*P* < 0.001. Black dots are mean values ± 1SE.

**Table 1.** Means (±SE) and statistical significance of mating type effects on life history traits measured in experimental lines of the hermaphrodite land snail *Rumina decollata*. Number of observations indicated in parentheses. *p < 0.05; **p < 0.01; ***p < 0.001. See Table S1 from Supp. Inf. for details on statistical models used.

| Traits measured | Mating treatment |  | Mating effect |
| --- | --- | --- | --- |
|  | Outcrossing | Selfing |  |
| Body weight | 1.74±0.04 (66) | 2.31±0.03 (45) | $F_{1,109} = 112.8^{***}$ |
| Age at first reproduction | 136.0±4.6 (35) | 193.0±6.9 (46) | $F_{1,79} = 41.5^{***}$ |
| Fecundity | 6.99±0.38 (68) | 2.87±0.46 (45) | $F_{1,111} = 47.6^{***}$ |
| Hatching time | 27.6±0.46 (62) | 31.8±1.82 (21) | $F_{1,81} = 10.4^{**}$ |
| Juvenile survival | 0.70±0.04 (68) | 0.30±0.06 (43) | $X^2_2 = 31.6^{***}$ |

### Individual growth

The growth parameters revealed distinct patterns across generations and mating treatments (Table 2, Figure 2A, B). In the F_1_, snails under *selfing* conditions exhibited a higher asymptotic shell length (*L∞* = 57.55 mm) compared to those under *outcrossing* conditions (*L∞* = 51.48 mm). The growth constant (*K*) was similar between treatments, with values of 2.23 ± 0.12 year⁻¹ for *selfing* and 2.18 ± 0.08 year⁻¹ for *outcrossing*. Both treatments displayed maximum seasonal oscillation (C = 1), with a similar starting point of oscillation (*ts* ≈ 2.16). The Growth Performance Index (GPI) revealed consistent differences between mating treatments being higher for the *selfing* treatment (3.87) compared to *outcrossing* (3.76), indicating faster growth under self-fertilization.

**Figure 2.**
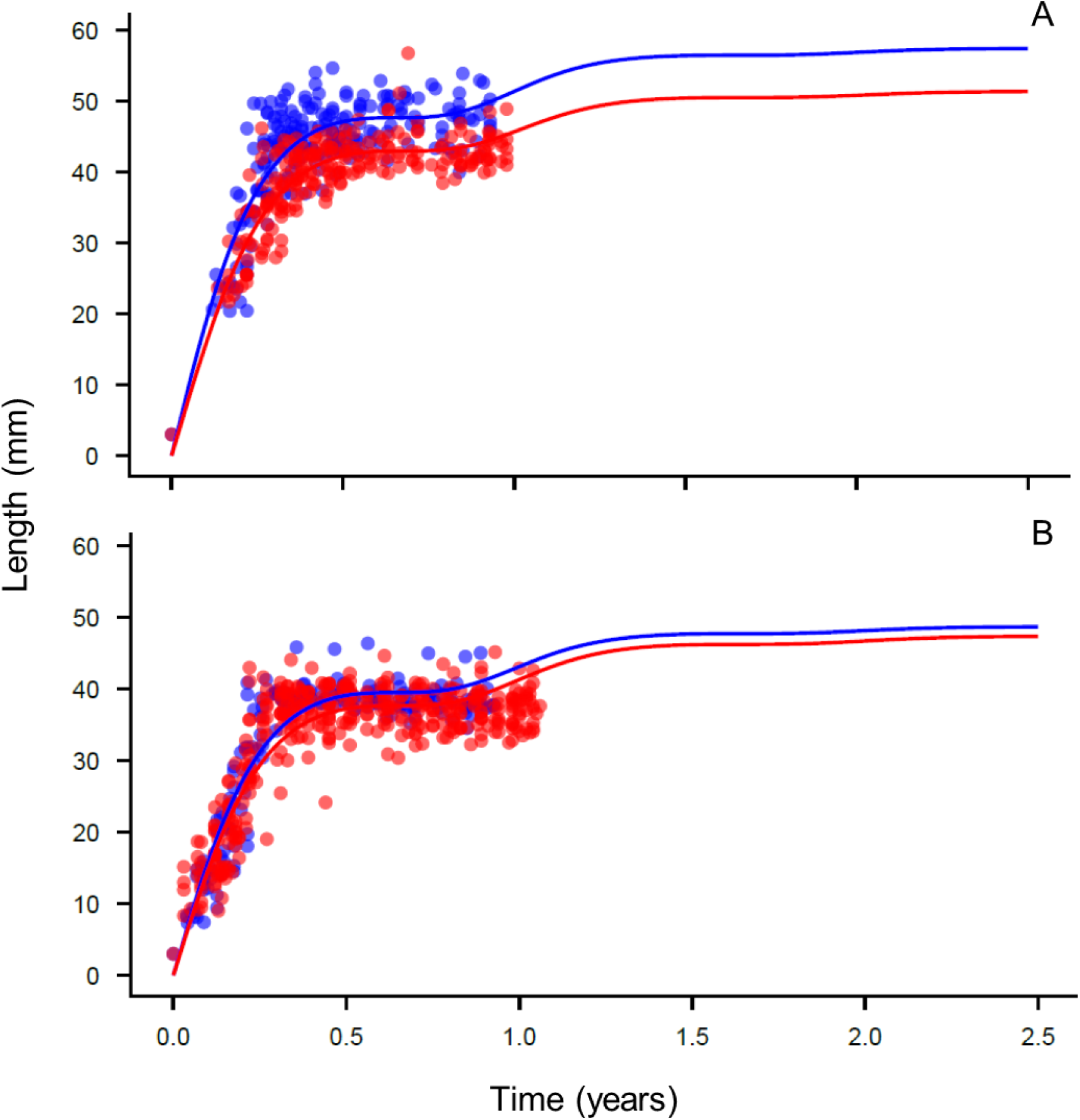
Growth curves fitted to the size-at-age data using the Specialized von Bertalanffy Growth Function (SVBGF) for the first (A) and second (B) generations of *Rumina decollata*. Red dots and lines represent the outcrossing treatment, while blue dots and lines represent the selfing treatment. Data points correspond to individual measurements, and the lines represent the model fit for each mating treatment.

**Table 2.** Growth parameters estimated using the seasonalized von Bertalanffy growth function (SVBGF) for two laboratory generations (first and second) and two mating treatments (outcrossing and selfing). L∞ represents the asymptotic shell length; *K* is the growth constant (year⁻¹) indicating the rate at which L∞ is approached; C represents the relative amplitude of seasonal oscillation (0 = no seasonal effect, 1 = maximum amplitude); and *ts* is the starting point of the oscillation. The Growth Performance Index (GPI) reflects the theoretical growth rate standardized to an organism with L∞ of 1 unit. Values are presented as mean ± standard error (SE).

| Growth parameters | First generation |  | Second generation |  |
| --- | --- | --- | --- | --- |
|  | Outcrossing | Selfing | Outcrossing | Selfing |
| $L_\infty$ | 51.48 | 57.55 | 47.53 | 48.82 |
| $K$ | 2.18 ± 0.08 | 2.23 ± 0.12 | 2.03 ± 0.06 | 2.15 ± 0.10 |
| $C$ | 1 | 1 | 1 | 1 |
| $ts$ | 2.18 ± 0.02 | 2.16 ± 0.02 | 2.15 ± 0.01 | 2.15 ± 0.02 |
| GPI | 3.76 | 3.87 | 3.66 | 3.71 |

In the F_2_, the *L∞* were smaller for both treatments compared to the first generation, with *outcrossing* at 47.53 mm and *selfing* at 48.82 mm. The *K* remained high but showed a slight decline relative to the first generation, with values of 2.03 ± 0.06 year⁻¹ for *outcrossing* and 2.15 ± 0.10 year⁻¹ for *selfing*. As observed in the first generation, the amplitude of seasonal oscillation remained maximal (C = 1) and the starting point of oscillation remained consistent (*ts* ≈ 2.15). The *GPI* followed a similar trend, with *selfing* (3.71) maintaining higher values compared to *outcrossing* (3.66).

### Shell morphometrics

The Procrustes superimposition and regression analysis revealed a significant allometric effect, with size explaining 21.23% of the shape variation (*P* <0.0001) in F_1_ snails from each treatment. To remove this effect, residuals from the regression were used for further analyses. The Principal Component Analysis (PCA) showed that the first two principal components (PC1 and PC2) accounted for 43.13% of the total shape variation (PC1 = 27.17%, PC2 = 15.96%). However, the Hotelling’s T-square test (T² = 19.4902, *P* = 0.9992 parametric, *P* = 0.9960 permutation) indicated no statistically significant differences in shell shape between snails from each mating treatment.

A similar pattern was observed in the F_2_ generation, with significant allometry detected (*P* < 0.0001), though size accounted for a lower proportion of shape variation (10.44%). The PCA revealed 35.40% of total variance explained by the first two principal components (PC1 = 20.78%, PC2 = 14.62%). The Hotelling’s T-square test for shape differences between treatments returned T² = 66.4771, with a parametric *P*-value of 0.2654 and a permutation *P*-value of 0.2710, indicating no significant morphological differentiation between the outcrossed and selfed groups in the second generation.

## Discussion

Our study provides the first comprehensive analysis of how different mating systems influence key life history traits, individual growth, and shell morphology in the hermaphroditic land snail *Rumina decollata*. By employing a controlled, multi-generational experimental design, this research goes beyond previous genetic and observational studies, shedding light on the physiological and ecological trade-offs associated with self-fertilization versus outcrossing. Our findings reveal clear patterns that deepen our understanding of reproductive strategies and their fitness consequences in *R*. *decollata*.

### Life history traits

The increased body weight at first reproduction in selfing snails, despite the lack of a significant effect of age at first reproduction as a covariate, suggests that other factors may be at play. Selfing snails may allocate more resources to somatic growth rather than reproduction during their delayed reproductive phase. The increased body weight at first reproduction in selfing snails could reflect a trade-off between growth and reproduction [21]. Physiological or developmental constraints of selfing may delay resource allocation to reproduction, leading to prolonged growth and larger body size before reproduction begins. This increased body size could serve as a compensatory strategy, enhancing reproductive output by providing more energy reserves for larger egg clutches, thus improving offspring survival if outcrossing occurs before the maximum waiting time is reached [6].

Selfing was associated with delayed reproduction, with snails laying their first clutch 39 days later than their outcrossing counterparts. While we did not explicitly assess the minimum age at maturity, previous studies report it to be approximately 10 weeks in laboratory conditions [16,17]. Our study used first-generation lab-reared snails, for which the hatching date was known. Snails in the outcrossing treatment were placed in mass mating at 12 weeks to ensure full maturity, which may imply that the age at first reproduction—and thus the waiting time—could be underestimated. Future research could benefit from protocols that monitor fecundity during the first week of reproductive life to provide a more accurate estimate of age at maturity (e.g., [6]). Interestingly, despite this caveat, the waiting time observed in this study (39 days) far exceeds the 16 days (see Table 1 from Selander et al. [17]). Nevertheless, it remains considerably shorter than waiting times reported for other terrestrial snails with mixed mating systems. For example, *Arianta arbustorum* and *Neohelix albolabris*, both with lifespans of 3–3.5 years, exhibit waiting times of 360–730 days and 100–270 days, respectively [44–46]. Similarly, *Partula taeniata*, which can live up to 10 years, delays reproduction by 605 days [10]. Even *Lissachatina fulica*, a highly invasive species and preferential outcrosser with a lifespan of 3–5 years, exhibits a waiting time of 240 days when isolated [47]. In contrast, the relatively short waiting time of *R*. *decollata* likely reflects its shorter lifespan (average of 1.5 years; Pizá, unpublished results) and underscores its efficiency as a strong colonizer, enabling rapid establishment in novel environments.

The mating treatment significantly influenced female fecundity, with selfing snails laying 51% fewer eggs per individual than their outcrossed. Additionally, eggs from the selfing treatment took, on average, 4 days longer to hatch than those from the outcrossing treatment. Similar reductions in fecundity have been observed in other land snail species. For instance, in *Arianta arbustorum*, isolated individuals exhibited a 50% decrease in fecundity compared to those reproducing via cross-fertilization [45]. One possible explanation for reduced fecundity in selfing snails is the absence of social facilitation—a phenomenon where interactions with conspecifics influence growth, metabolism, behavior, and reproduction [48]. This has been documented in pulmonate gastropods, where exposure to conspecific mucus increases clutch size and egg production, likely due to stimulatory substances in the mucus [49]. Similarly, copulation and spermatophore transfer can enhance oviposition [9,50]. In the obligate selfer *Balea perversa*, pair-reared individuals produced more eggs over extended periods compared to isolated individuals, suggesting that social facilitation can shape life-history traits [7]. Although social facilitation could partly explain the reduced fecundity observed in the selfing treatment, it is unlikely to account for a 51% reduction. To minimize its potential effects, snails in this study were isolated in individual boxes after mass mating, and box lids were alternated weekly within each treatment to homogenize any residual cues. The consistent and substantial reduction in fecundity is more plausibly attributed to inbreeding depression, which is known to affect reproductive output and offspring viability in selfing populations [12]. The delay in hatching time for eggs from the selfing treatment also suggests developmental differences that may reflect physiological constraints or reduced embryonic viability. Prolonged incubation periods could be a manifestation of inbreeding depression, as genetic homozygosity may lead to slower development or weaker embryos [11]. This aligns with our observation of reduced juvenile survival in the selfing treatment.

We observed a significant decrease in juvenile survival in the selfing treatment compared to the outcrossing treatment. Juvenile survival is a critical component of fitness, as it manifests well before reproduction begins and affects both male and female individuals. Furthermore, inbreeding depression tends to be most pronounced during this stage of the life cycle [27,28]. The apparent inbreeding depression (AID), measured as the reduction in juvenile survival in the selfing treatment, was 0.57. This value indicates notable inbreeding depression, similar to that observed in other widespread species such as the freshwater snail *Physa acuta* [10]. However, when compared to other land snails that are known preferential outcrossers, the AID in *Rumina decollata* reflects an intermediate level of inbreeding depression, as values for other species are nearly twice as high (e.g., *Succinea putris* = 0.97, *Partula taeniata* = 0.98, *Neohelix albolabris* = 0.96, *Arianta arbustorum* = 0.37; Escobar et al [10] and references therein). These findings support the idea that high outcrossing rates are associated with elevated inbreeding depression and longer waiting times for reproduction [6,10].

Interestingly, another way to interpret this result is that 32% of selfing snails in our study produced viable offspring, which is markedly higher than one of the most invasive land snails, *Lissachatina fulica*. In a common garden laboratory experiment, of 116 isolated individuals of this widespread pest species, only 8 reproduced via self-fertilization (7%), and of those, only 13–40% laid a viable clutch, yielding an overall viability of 0.9–3% [47]. Even the highest viability estimate for *L*. *fulica* remains 11 times lower than what was observed for *R*. *decollata*.

The high reproductive capacity of *R*. *decollata* under strong ecological constraints, such as low population density or mate absence, underscores its ability to persist and reproduce in challenging environments. This facultative self-fertilization strategy, combined with its extensive geographic range across diverse biogeographic and climatic conditions [15], suggests that *R*. *decollata* has the potential for further population expansion, making it a species of growing ecological concern.

### Individual growth

Individual growth rate was different between mating treatments and across generations. As F_1_ snails from both treatments were descendants of wild outcrossing parents, the observed differences are likely a result of experimental conditions—specifically, the isolation of the selfing group versus the mass mating of the outcrossing group. Hence, the higher growth performance (GPI) observed in the selfing treatment for F_1_ snails suggests that physiological or developmental constraints associated with selfing may delay resource allocation to reproduction. This delay may allowed prolonged growth and a larger shell size before reproduction begins, potentially reflecting a growth-reproduction trade-off [21]. Although selfing confers short-term advantages in growth, these benefits may come at a cost to long-term fitness, as inbreeding depression could limit subsequent generational performance. Outcrossing, by maintaining genetic diversity, avoids these immediate physiological advantages but ensures greater resilience over time. These findings align with broader reproductive strategy patterns, where environmental conditions and life-history traits shape the balance between short-term and long-term benefits [51].

The reduction in GPI from F_1_ to F_2_ in both treatments indicates generational effects on growth. For the selfing treatment, the sharper decline in GPI can be attributed to inbreeding depression, resulting from the expression of deleterious recessive alleles due to reduced heterozygosity [11]. This emphasizes the cumulative genetic costs of self-fertilization despite the initial growth advantages observed in F_1_. In contrast, in the outcrossing treatment, the decline in GPI was less pronounced, likely reflecting the benefits of preserved genetic diversity. However, even in outcrossing populations, factors such as genetic drift or subtle genetic constraints could contribute to the observed decline. For instance, if a small subset of individuals disproportionately contributed to the F_2_ population, this could inadvertently reduce the effective genetic diversity despite controlled mass mating [52]. The lower GPI observed in F_2_ individuals across both treatments may also reflect physiological or ecological constraints on growth potential in subsequent generations. Studies on mollusks and other invertebrates suggest that multigenerational exposure to controlled environments can impose cumulative genetic or epigenetic effects, limiting growth and fitness improvements over time [53]. This decline may also result from subtle environmental stressors or near-optimal growth conditions in F_1_, leaving limited scope for further enhancements in F_2_. However, the scope of our study, limited to two laboratory generations, is insufficient to confirm such processes conclusively. Future research involving additional generations and natural conditions would be required to robustly assess cumulative genetic or epigenetic constraints.

### Shell Morphometrics

The lack of statistically significant shape differences between selfed and outcrossed snails in both generations suggests that variations in GPI did not strongly translate into morphological divergence in shell shape. Although previous studies have reported associations between growth rate and shell shape variation (e.g., [22,23]), our results indicate that selfing and outcrossing may influence growth without necessarily inducing distinct morphological differentiation in *Rumina decollata*. Further studies incorporating additional generations could help clarify whether subtle morphological changes emerge over longer timescales.

### Environmental, Economic, and Health Impact

Beyond its ecological and reproductive traits, *R*. *decollata* poses potential environmental, economic, and health risks, warranting close monitoring as a potentially invasive species. Recognizing its impact, *R*. *decollata* was recently added to the “Official List of Invasive Alien Species of Argentina” as a “Restricted Species”, indicating its high environmental and socioeconomic impact [54]. Given its successful establishment and expansion across multiple regions, its ongoing spread may have unforeseen ecological consequences, particularly in non-native habitats where it may compete with or displace native gastropod species. Observations indicate a negative impact of *R*. *decollata* on the native land snail *Bulimulus bonariensis* [15], suggesting potential threats to local biodiversity. Additionally, *R*. *decollata* has been identified as a potential intermediate host for cat-parasitic nematodes, including *Aelurostrongylus abstrusus* and *Toxocara cati* [55–57]. As urban populations of *R*. *decollata* increase, particularly in areas with high densities of domestic and stray cats, the risk of parasite transmission may also rise. Since these nematodes cause respiratory diseases in felines, and their spread is facilitated by the presence of gastropod intermediate hosts, *R*. *decollata*’s proliferation in urban environments could increase disease incidence and pose indirect health risks to companion animals and potentially human populations. These ecological and health concerns emphasize the need for ongoing surveillance and management strategies to mitigate the potential impact of this alien species as it continues expanding into new habitats.

## Conclusion

Our findings highlight the significant fitness costs associated with self-fertilization in *Rumina decollata*, evidenced by delayed reproduction, reduced fecundity, and lower juvenile survival. These effects are consistent with pronounced inbreeding depression. Outcrossing, in contrast, provides clear advantages by sustaining reproductive success and life history traits across generations. Growth and shell morphometric patterns further reveal distinct energy allocation strategies between selfing and outcrossing treatments. By highlighting the nuanced interplay between mating systems, this study enhances our understanding of *R*. *decollata*’s reproductive strategies in shaping life history traits under environmental constraints. As this land snail continues its marked geographic expansion across varying latitudes, these insights emphasize its capacity as an alien species to thrive and colonize diverse and often unpredictable habitats.

**Supplementary Materials:** The following supporting information can be downloaded at:

**Supplementary Methods and Results**: Geometric apex and shell length measurement

**Supplementary Tables:** Table S1: Detailed results on linear models

**Supplementary Figures:** Figure S1: Breeding protocol; Figure S2: Geometric apex of the decollate shell of *Rumina decollata*.

## Author Contributions

Conceptualization, resources, validation, writing—review and editing, supervision and funding acquisition, J.P and N.B.; writing—original draft preparation, J.P, L.C and N.B.; methodology, N.B; investigation and data curation, J.P., L.C., M.P., M.A., and N.B.; formal analysis, L.C, M.A. and N.B.. All authors have read and agreed to the published version of the manuscript.

## Institutional Review Board Statement

Not applicable.

## Informed Consent Statement

Not applicable. All procedures contributing to this work comply with the ethical standards of the relevant national and institutional guides.

## Data Availability Statement

The data that support the findings of this study will be made available in the Dryad Digital Repository upon publication of this paper: https://datadryad.org/stash.

## Acknowledgments

JP and NB are staff researchers at CONICET (Consejo Nacional de Investigaciones Científicas y Técnicas, Argentina). LC, MP, and MA were undergraduate students in the Becas de Estímulo a las Vocaciones Científicas (EVC CIN) program. We also wish to thank the reviewers for their valuable feedback and suggestions during the peer-review process.

