## Supplemental material for "Impact of mating strategies on life-history traits in the alien land snail *Rumina decollata*"

**<u>Table of Contents</u>**

***Supplementary Methods***
Reconstruction of geometric apex 1
References 1

***Supplementary Tables***
*Table S1*. Results of linear models for life history traits 2

***Supplementary Figures***
*Figure S1*. Breeding protocol 3
*Figure S2*. Reconstruction of geometric apex 4

### Supplementary Methods

#### *Reconstruction of geometric apex*

We measured the shell length using the ImageJ software on each image. To reconstruct the missing apex lost during decollation, we drew projection lines tangent to the spire, connecting the sutures and extending them to their intersection, which marks the geometric apex (Figure. S2). While the reconstructed shell lengths may not reflect the actual lengths, this method enables a systematic study of individual growth, accounting for the lost whorls and the energy invested in their formation [1, 2].

### Supplementary Tables

**Table S1.** Detailed results of the linear models on fitness traits measured on $F_1$ and $F_2$ offspring resulting from different mating treatments (selfing and outcrossing) of the hermaphrodite snail *Rumina decollata*. Significant values of each effect are indicated in bold.

| Variable | Fixed effects | | | Random effects | Number of observations | Model |
|---|---|---|---|---|---|---|
| | Mating treatment | Age at first clutch | Weight of focal | Individual | | |
| Body weight | $F_{1,81}$= 56.7 | | | | 83 | Gaussian |
| | *P* **<0.001** | | | | | |
| (Age at first clutch as covariable) | $F_{1,52}$ = 22.43 | $F_{1,52}$= 1.08 | | | 55 | |
| | *P* **<0.001** | *P* = 0.304 | | | | |
| Age at first clutch | $F_{1,55}$ = 7.3 | | | | 57 | quasi-Poisson |
| | *P* **= 0.009** | | | | | |
| Fecundity | $F_{1,83}$ = 148.9 | | | | 85 | quasi-Poisson |
| | *P* **<0.001** | | | | | |
| (Weight of focal as covariable) | $F_{1,83}$ = 24.16 | | $F_{1,83}$ = 0.12 | | 85 | |
| | *P* **<0.001** | | *P* = 0.727 | | | |
| Hatching time | $F_{1,53}$ = 2.2 | | | | 55 | Poisson |
| | *P* **= 0.028** | | | | | |
| Juvenile survival | $X^2_2$ = 25.7 | | | variance = 4.95 | 83 | Binomial |
| | *P* **<0.001** | | | *P* **<0.001** | | |

### Supplementary Figures

**Figure S1.** Breeding protocol for the experimental populations. Diagram depicting the experimental setup for the two mating treatments: (A) Outcrossing treatment, where virgin snails (n = 70) were kept in isolation until fully mature, then placed in a terrarium for one week of mass mating to promote cross-fertilization, followed by re-isolation to lay eggs. (B) Selfing treatment, where mature snails were kept in isolation until they laid eggs, enforcing obligate self-fertilization. The offspring from each clutch (Group A and B) are full-sibs, and one juvenile per clutch (focal snail) was isolated and raised until maturity to form two independently maintained laboratory populations for each treatment.

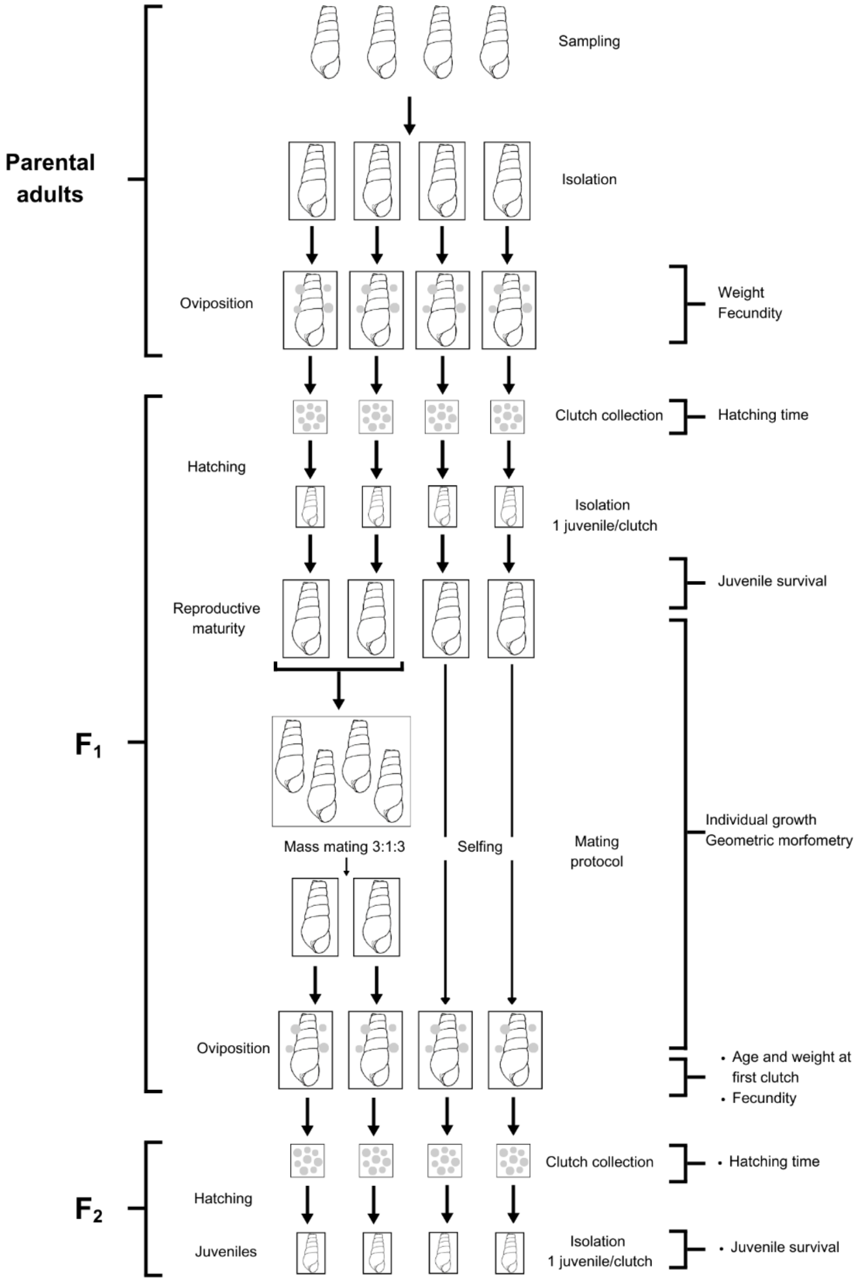

**Figure S2.** Reconstruction of the geometric apex. Diagram illustrating the method used to reconstruct the missing apex of decollated shells. Projection lines are drawn tangent to the spire, connecting the sutures and extending them until their intersection, which defines the geometric apex. This method allows for a systematic study of individual growth by accounting for the lost whorls and the energy invested in their formation.

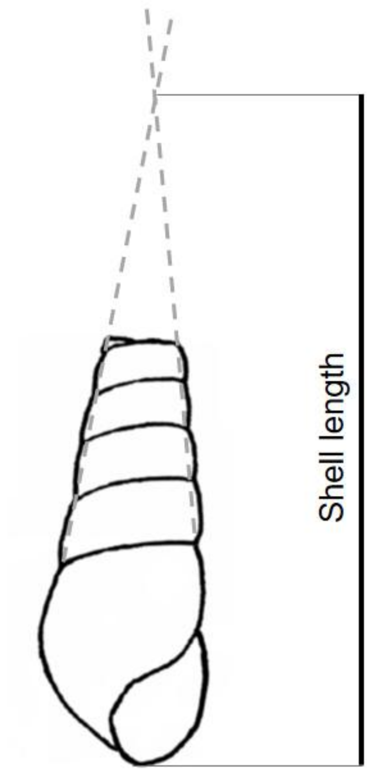
